# Tirzepatide improves pancreatic β-cell function in mice and patients with type 2 diabetes

**DOI:** 10.64898/2026.02.13.704343

**Authors:** Zhehui Li, Jiacheng Guo, Ying Cheng, Tongyu Zhang, Xiumei Luo, Shuyi Zhang, Qi Ren, Zhiqiang Wu, Ning Chen, Mingyu Li

**Affiliations:** State Key Laboratory of Cellular Stress Biology and Fujian Provincial Key Laboratory of Innovative Drug Target Research, School of Pharmaceutical Sciences and School of Life Sciences, Xiamen University, Xiamen 361102, China; Department of Endocrinology, Zhongshan Hospital (Xiamen), Fudan University, Xiamen 361000, China; School of Marine Life Science, Ocean University of China, Qingdao 266003, China; State Key Laboratory of Vaccines for Infectious Diseases, Xiang An Biomedicine Laboratory, Xiamen University, Xiamen 361102, China; School of Medicine, Xiamen University, Xiamen 361102, China

**Keywords:** Tirzepatide, Type 2 diabetes, dual GIP/GLP-1 receptor co-agonist, β-cell identity, FOXO1

## Abstract

The dual incretin receptor agonist tirzepatide improves β-cell function in T2D patients, but the underlying mechanism remains unclear. This study aimed to elucidate the molecular pathway through which tirzepatide restores β-cell functional improvement. High-fat diet (HFD)-fed C57BL/6J mice were treated with vehicle, a GIP analogue, semaglutide or tirzepatide. Tirzepatide significantly reduced body weight and improved glucose tolerance in HFD-fed mice without altering β-cell mass, proliferation, or apoptosis. Instead, tirzepatide reversed β-cell dedifferentiation, as indicated by reduced ALDH1A3 expression and restored levels of the identity transcription factors PDX1 and MAFA. Single-cell RNA sequencing (scRNA seq) and *in vitro* studies revealed that tirzepatide up-regulated FOXO1, reactivating the FOXO1-PDX1/MAFA axis. In T2D patients, tirzepatide improved glycemic control, reduced insulin demand, increased HOMA-β, and decreased HOMA-IR. Improvement in HOMA-β correlated positively with baseline insulin resistance. Hence, our study suggested that tirzepatide restores β-cell function in T2D by reprogramming stressed β cells and re-establishing β-cell identity through FOXO1-dependent transcriptional reactivation. These findings provide a mechanistic basis for the superior efficacy of dual incretin receptor agonism in T2D management.

**ARTICLE HIGHLIGHTS:**

- Tirzepatide restores β cell identity and function without altering β cell mass in HFD induced diabetic mice.
- Tirzepatide reverses β-cell dedifferentiation and restores key β-cell transcription factors (PDX1, MAFA) through reactivation of the AKT-FOXO1 signaling pathway.
- Tirzepatide increases HOMA-β and decreases HOMA-IR in T2D patients, and improvements in HOMA-β positively correlate with baseline insulin resistance.
- These results demonstrate that tirzepatide’s therapeutic benefits are not only metabolic but also involve direct restoration of β-cell identity and function. This highlights β-cell reprogramming as a novel therapeutic avenue, thus supporting the broader clinical adoption of dual incretin receptor agonists.

## 1. INTRODUCTION

Type□2 diabetes mellitus (T2D) is a complex metabolic disorder characterized by chronic hyperglycemia, insulin resistance and progressive pancreatic β-cell dysfunction [1, 2]. In pre- or early type 2 diabetes stage, compensatory hyperinsulinemia maintains euglycemia [3]. During the development of the diabetes, chronic metabolic stress, glucolipotoxicity, and inflammation ultimately drive β-cell decompensation and β-cell failure [4, 5]. Although loss of β-cell mass or ratio is the factor for β-cell failure, loss of β-cell identity and β-cell dedifferentiation also play a central role to β-cell failure, which accompanied by downregulation of key transcription factors such as PDX1 and MAFA [6, 7]. Moreover, Obesity-driven insulin resistance further exacerbates metabolic stress through ectopic lipid accumulation and adipose inflammation, and led to β-cell dysfunction [8].

Glucagon-like peptide-1 (GLP-1) and glucose-dependent insulinotropic polypeptide (GIP) are two main incretin hormones, and they are critical for postprandial glucose homeostasis, insulin secretion, lipid (fat) metabolism, and energy expenditure [9]. Due to the glucoregulatory and anorectic activities of GIP and GLP-1, the incretin-based therapies, GLP-1 agonists or GIP and GLP-1 co-agonists, have been successfully developed for treatment of type 2 diabetes and obesity [10]. Semaglutide mimics the action of the incretin hormone GLP-1, which binds to GLP-1 receptors in pancreatic β-cells and extra pancreas. Semaglutide enhances insulin secretion in a glucose-dependent manner, suppresses glucagon release, delays gastric emptying, and promotes satiety, thereby benefit for therapy of type 2 diabetes and obesity [11, 12]. Tirzepatide is a dual GLP-1/GIP receptor agonist, which synergistically enhances β-cell function, shown remarkable efficacy in lowering HbA1c and body weight in patients with type 2 diabetes and patients with obesity [13, 14]. Moreover, tirzepatide displayed better reductions in HbA1c and more substantial weight loss compared to semaglutide [15]. Furthermore, tirzepatide significantly improved β-cell function and insulin sensitivity compared with semaglutide [16]. However, the molecular mechanisms underlying tirzepatide induced β cell improvement remain incompletely understood.

In this study, by using a high-fat diet (HFD) induced T2DM mouse model and clinical patient data, we revealed that tirzepatide not only ameliorates systemic metabolic dysfunction but also reprograms stressed β-cells to restore transcriptional identity and functional competence, without expanding β-cell mass. By integrating physiological, histological, molecular, and single-cell transcriptomic analyses, we uncover a critical role for the FOXO1-PDX1/MAFA axis in mediating tirzepatide’s restorative effects on β-cell function.

## 2. RESEARCH DESIGN AND METHODS

### 2.1 Animals

Male C57BL/6J mice were purchased from GemPharmatech (Nanjing, China). The mice were maintained in a specific pathogen-free environment at the Xiamen University Animal Center. They were kept in individually ventilated cages with no more than five mice per cage, under a 12-hour light/dark cycle at a controlled temperature of 20-23 □, and had free access to standard chow and water. All mice were randomly assigned to normal chow-fed (Control) or high-fat diet (HFD) groups. The HFD group was gradually transitioned to a 60% HFD (low-fat: high-fat diet ratios of 9:1, 7:3, 5:5, 3:7, and 1:9 every two days). After complete transition, mice were maintained on a 60% HFD for 12 weeks to induce obesity [17]. Only male mice were used in this study, and all experimental procedures were conducted in accordance with protocols approved by the Institutional Animal Care and Use Committee (Animal Ethics Approval: FDULAC20250381) and complied with Fudan University Laboratory Animal Center guidelines.

### 2.2 Drug Administration

Tirzepatide (LY3298176; Sellback), semaglutide (Ozempic, Novo Nordisk), and [D-Ala²]-GIP (Nanjing Peptide Valley) were used. Tirzepatide was initially prepared as a 100 mg/mL clear stock solution in DMSO, and subsequently diluted by mixing 50μL of the stock solution with 400 μL polyethylene glycol 300 (PEG300) until the solution became clear, followed by the addition of 50 μL Tween 80 (T8220; Beijing Solarbio Science & Technology Co., Ltd) with thorough mixing. Finally, ddH2O was added to a final volume of 1mL. The working solution was freshly prepared before use. Tirzepatide and [D-Ala²]-GIP were dissolved in 40 mM Tris-HCl, whereas semaglutide was dissolved in 0.9% NaCl. During the 12 weeks of dietary induction, body weight and fasting blood glucose were recorded weekly. Control mice received vehicle (40 mM Tris-HCl), while HFD-fed mice were randomly allocated to receive daily subcutaneous injections for 2 weeks of: vehicle, [D-Ala²]-GIP (10 nM), semaglutide (10 nM), or tirzepatide (3 nM or 10 nM) (*n* = 5 per group).

### 2.3 Intraperitoneal Glucose Tolerance Test (IPGTT)

After 2 weeks of treatment, mice were fasted for 12 h with free access to water before IPGTT. Fasting blood glucose (0 min) was measured from the tail vein using a glucometer (Yuwell, China). Mice then received an intraperitoneal injection of 20% glucose (0.1 mL/10 g body weight). Blood glucose was measured at 15, 30, 60, 90, and 120 min post-injection.

### 2.4 Blood and Tissue Collection

Mice were fasted overnight for 12 h before harvested. Blood was collected via orbital bleeding, allowed to clot for 30 min at room temperature, and centrifuged at 3000 × g for 10 min at 4 °C. Serum was stored at -80 °C. Pancreas and liver were fixed in 4% paraformaldehyde, cryoprotected in sucrose, and embedded in OCT. White adipose tissues (inguinal, epididymal) and brown adipose tissue (interscapular) were dissected, weighed, and photographed.

### 2.5 Biochemical and Hormonal Assays

Serum levels of alanine aminotransferase (ALT, B1008), aspartate aminotransferase (AST, B1058), high-density lipoprotein (HDL, B1003), low-density lipoprotein (LDL, B1004), total cholesterol (T-CHO, B1067), and triglycerides (TG, B1066) were measured using commercial kits (Nanjing Jiancheng Bioengineering Institute). Serum insulin (MM-0579M1), glucagon (MM-0167M1), and total GLP-1 (MM-0027M1) were quantified using ELISA kits (Jiangsu Meimian Industrial Co., Ltd.) according to the manufacturers’ protocols.

### 2.6 Immunofluorescence Staining

Frozen pancreatic sections (10 μm) were prepared using a Leica cryostat. Sections were permeabilized with 0.3% Triton X-100 for 3 h, blocked with 5% FBS in PBS containing 0.1% Tween 20 for 1 h, and incubated overnight at 4 °C with primary antibodies (anti-glucagon, Abcam ab92517; anti-insulin, Cell Signaling Technology, L6B10; anti-insulin, Biosynth, 20-IP35; anti-somatostatin, Abcam, ab30788; anti-ALDH1A3, Novus Biologicals, NBP2-15339; anti-PDX1, Abcam, Ab219207; anti-MAFA, Abcam, Ab17976; anti-FOXO1, Proteintech, 18592-1-AP; anti-p-FOXO1(Ser256), Affinity, AF3417), followed by fluorescent secondary antibodies for 2 h at room temperature. Whole-pancreas sections were scanned using an Olympus VS200 slide scanner, while individual islets were imaged using a Leica SP8 confocal microscope. ImageJ (v1.54f) was used for quantification of islet number, endocrine cell area, and fluorescence intensity.

### 2.7 Islet Isolation and Single-Cell Suspension

Pancreatic islets were isolated after perfusion *via* the common bile duct with 3 mL collagenase type IV (1 mg/mL, Gibco, 17104019). Digested tissue was incubated at 37 °C for 10 min, and islets were purified by density gradient centrifugation and hand-picked under a stereomicroscope. Islets were dissociated into single cells using trypsin at 37 °C for 15 min and filtered through a cell strainer. Viability (>80%) and cell counts were determined by trypan blue exclusion.

### 2.8 Single-Cell RNA Sequencing

Single-cell RNA sequencing (scRNA-seq) of pancreatic islets from three mice per group (chow-fed, HFD-fed, and HFD-fed with 10 nM Tirzepatide) was performed by a commercial service provider (Illumina NovaSeq X Plus). Raw sequencing data were processed using Seurat (v4.3.2) [18]. Low-quality cells were excluded if they had >25% mitochondrial genes, <200 or >7,500 detected genes. The remaining data were normalized using the *LogNormalize()* method, and highly variable genes were identified with *FindVariableFeatures()*. Principal component analysis (PCA) was performed, and significant principal components were used for clustering (*FindClusters()*) and marker identification (*FindAllMarkers()*). Dimensionality reduction was conducted using UMAP and t-SNE. Cell types were annotated according to canonical marker genes, resulting in ten major pancreatic cell types, including β cells, α cells, δ cells, endothelial cells, immune cells, PP cells, stellate cells, mesenchymal cells, acinar cells, and ductal cells. The relative proportions of each cell type were calculated and visualized using the *dplyr* package. For β-cell specific analysis, β cells were extracted as a subset, and gene expression patterns were visualized using violin plots (*VlnPlot()*). Differentially expressed genes (DEGs) were identified with the *limma* package [19], and KEGG pathway enrichment was performed using the *enrichKEGG()* function [20].

### 2.9 Machine Learning-Based Gene Selection

To further refine key DEGs, three complementary machine learning algorithms were applied, including LASSO regression (*glmnet* package), Boruta feature selection (*Boruta* package), and Extreme Gradient Boosting (XGBoost; *xgboost* package). For LASSO, the *cv.glmnet(*) function was used to perform 10-fold cross-validation and identify the optimal λ parameter in a penalized logistic regression model [21]. Boruta analysis employed *Boruta()* followed by *TentativeRoughFix()* to confirm important genes [22]. For XGBoost, the *xgb.train()* function was run with a binary logistic objective and 100 boosting iterations to compute gene importance scores (*xgb.importance()*) [23]. Genes identified by at least two of the three algorithms were defined as key candidate genes for downstream pathway enrichment and functional analyses.

### 2.10 Cell Culture, cell line treatment and Western Blotting

Min6 cell was purchased from the American Type Culture Collection (ATCC; Manassas, VA, USA) and maintained in high-glucose DMEM (Gibco, 11965092) supplemented with 10% fetal bovine serum (FBS) and 1% penicillin/streptomycin at 37 °C in a humidified incubator with 5% CO_2_. PA (Sigma Aldrich P9767) was diluted in water at 65 °C to a concentration of 100 mmol/L and finally diluted in serum free Roswell Park Memorial Institute (RPMI) 1640 containing 10% bovine serum albumin (BSA) (Sigma Aldrich A730) and 25 mmol/L glucose to a concentration of 300 μM. Cells were divided into the control group, HG + PA group (25 mmol/L glucose + 300 μmol/L PA), and HG + PA + TZP (25 mmol/L glucose + 300 μmol/L PA + 10 nM Tirzepatide). The cells were treated for 24 hours before further experiments.

After the treatment, cells were lysed in radioimmunoprecipitation assay (RIPA) buffer, and protein concentrations were determined using a BCA Protein Assay Kit (Nanjing Vazyme Biotech Co., Ltd, E112-01). Equal amounts of protein were separated by SDS-PAGE and transferred to PVDF membranes. After blocking in 5% nonfat milk in TBST, membranes were incubated overnight at 4 °C with primary antibodies, followed by horseradish peroxidase-conjugated secondary antibodies for 1 h at room temperature. Bands were visualized using enhanced chemiluminescence (ECL) and imaged with a Bio-Rad chemiluminescent detection system. Primary antibodies against GAPDH, AKT1, p-AKT(Ser473), FOXO1 were from Proteintech (GAPDH, 60004-1-Ig; AKT1, 10176-2-AP, p-AKT (Ser473), 66444-1-Ig, FOXO1, 18592-1-AP); antibodies against p-FOXO1(Ser256), PDX1, MAFA, and ALDH1A3 were from Affinity (AF3417), Abcam (Ab219207), Abcam (Ab17976) and Novus Biologicals (NBP2-15339), respectively. HRP-conjugated secondary antibodies were obtained from Yeasen.

### 2.11 Clinical Study Population and Design

Fourteen adults with T2DM from February 2025 to November 2025 were recruited from the Department of Endocrinology, Zhongshan Hospital, Fudan University (Xiamen). Eligible participants were 18-70 years old, of any sex, with a disease duration of ≤5 years, a body mass index (BMI) of 24-40 kg/m², and glycated hemoglobin (HbA1c) between 6.5% and 9.0%. All patients met the 2018 American Diabetes Association’s (ADA’s) criteria for the diagnosis and classification of diabetes. None had received glucose-lowering therapy within six months prior to enrollment [24]. All participants provided written informed consent, and the study was approved by the Ethics Committee of Zhongshan Hospital (Xiamen), Fudan University (No. B2024-123R)

Exclusion criteria included type 1 diabetes, gestational diabetes, or other specific forms of diabetes; recent severe diabetic complications; acute or chronic pancreatitis; personal or family history of medullary thyroid carcinoma or multiple endocrine neoplasia type 2; significant hepatic or renal impairment (ALT or AST >2.5× upper limit of normal or eGFR <60 mL/min/1.73 m²); severe psychiatric or neurological disorders; alcohol or drug abuse; pregnancy or lactation; malignant tumor; known hypersensitivity to Tirzepatide; or use of any medications affecting body weight or islet β-cell function. Participants deemed by investigators to have poor compliance or otherwise unsuitable for study participation were also excluded.

Tirzepatide was administered at 2.5 mg once weekly and increased by 2.5 mg every four weeks if the patient tolerated. Participants who experienced significant adverse effects, primarily gastrointestinal symptoms, and declined dose escalation continued at the prior dose. Treatment lasted 12 weeks. Anthropometric and metabolic parameters, including body weight, height, fasting plasma glucose (FPG), fasting insulin, HbA1c, fasting C-peptide, liver and renal function, and lipid profiles, were measured before and after treatment according to standardized clinical laboratory operating procedures (SOPs). Insulin resistance and β-cell function were assessed using HOMA-IR and HOMA-β.

Of the 14 participants enrolled, 13 completed the study, with one participant withdrawing after four weeks due to severe constipation.

### 2.12 Correlation Analysis of Clinical Parameters in Patients

All statistical analyses were conducted in R. Pearson correlations among clinical and metabolic variables were computed using *rcorr()*, and the resulting correlation coefficients and p-values were visualized with *corrplot()* and chord diagrams (*circlize* package) to display strength, direction, and significance[25]. Univariate relationships between ΔHOMA□β and individual predictors were explored using scatter plots with regression lines and 95% confidence intervals.

### 2.13 Statistical Analysis

Data were analyzed with GraphPad Prism 8 (GraphPad Software v8.3.0, La Jolla, CA, USA) and presented as mean ± SEM. For normally distributed data, one-way ANOVA with Bonferroni post hoc tests was applied. For non-normally distributed data, paired Student’s t-test (one group), Mann-Whitney U test (two groups) or Kruskal-Wallis test with Dunn’s post hoc test (three or more groups) were used. *p* < 0.05 was considered statistically significant.

## 3. RESULTS

### 3.1 Tirzepatide reduces body weight and improves glucose tolerance in HFD-fed mice

Mice were subjected to high-fat diet (HFD) feeding with 12 weeks and followed with different drugs administration for 2 weeks, including vehicle, 10 nM [D-Ala2]-GIP, 10 nM semaglutide, 3 nM tirzepatide, and10 nM tirzepatide (Fig.1A). As shown in Fig. 1B-1C, 10 nM semaglutide, 3 nM tirzepatide and 10 nM tirzepatide significantly reduced body weight and suppressed food intake in HFD-fed mice, but not the 10 nM [D-Ala2]-GIP (Fig. 1B-1C and Fig. S1). In terms of weight loss efficacy, 10 nM tirzepatide was superior to 10 nM semaglutide, which in turn outperformed 3 nM tirzepatide (Fig. 1B-1C). IPGTT test revealed that 10 nM semaglutide, 3 nM tirzepatide, and 10 nM tirzepatide all improved glucose tolerance in HFD-fed mice, with 10 nM tirzepatide had the best therapeutic effect (Fig. 1D-1E).

**Figure 1.**
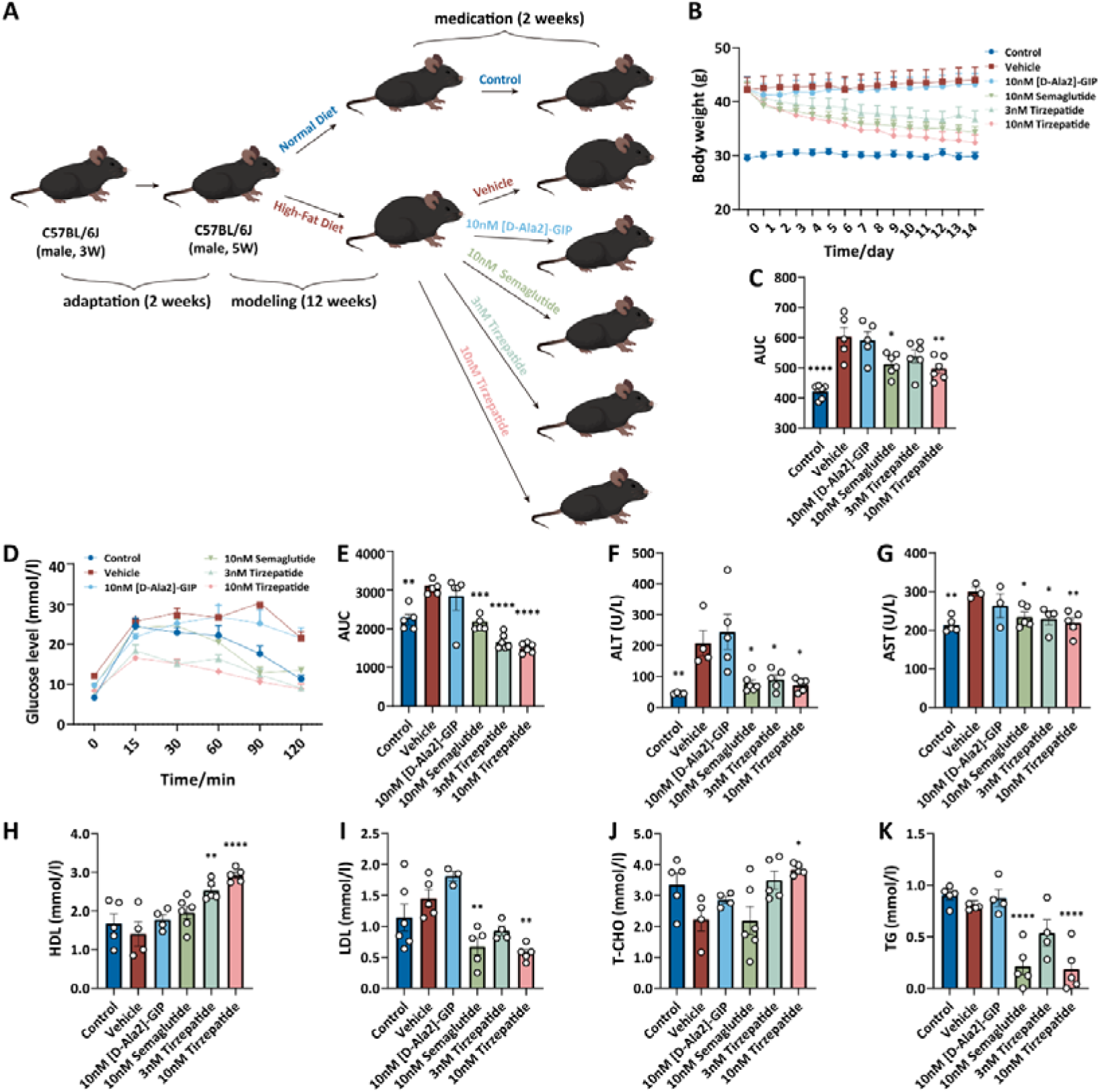
Tirzepatide reduces body weight and increases glucose tolerance in HFD-fed mice. **(A**) Schematic illustration of mouse modeling and drug treatment procedure. (**B-C**) The dynamics of body weight (**B**) and its AUC analysis (**C**) during drug treatment, mice n ≥ 5. (**D-E**) Intraperitoneal glucose tolerance test (IPGTT) (**D**) and its area under the curve (AUC) (**E**) of different groups. mice n ≥ 5. (**F**-**K**) Serum levels of ALT (**H**), AST (**I**), HDL (**J**), LDL (**K**), T-CHO (**L**) and TG (**M**) in each group, mice n ≥ 3. Data are presented as mean ± SEM, with significance determined by one-way ANOVA compared to the Vehicle group: \**p* < 0.05, \*\**p* < 0.01, \*\*\**p* < 0.001 and \*\*\*\**p* < 0.0001.

Moreover, 10 nM semaglutide, 3 nM tirzepatide, and 10 nM tirzepatide reduced the liver weight of HFD-fed mice (Fig. S2A). For adipose masses, these drug treatments also had trend to reduce eWAT, iBAT, and iWAT, except for 3 nM tirzepatide treated iWAT (Fig. S2B-S2D). Consistently, tirzepatide improved circulating biochemical parameters, including ALT, AST, HDL, LDL, T-CHO and TG (Fig. 1F-1K), indicating a systemic improvement in hepatic and lipid metabolism. In contrast, mice treated with 10 nM [D Ala2] GIP showed no significant differences from the HFD group across all measured biochemical parameters. Among all treatment groups, mice treated with 10 nM [D-Ala2]-GIP showed no significant differences from the HFD group across all measured circulating biochemical parameters. Notably, 10 nM tirzepatide produced the most pronounced therapeutic outcomes, supporting its superior potency. Together, these data demonstrated that 10 nM tirzepatide effectively reverses HFD-induced obesity and metabolic imbalance, improves both energy expenditure and insulin sensitivity.

### 3.2 Tirzepatide restores ***β***-cell function in HFD-fed mice

We next to investigated whether these drug treatments improve the pancreatic β-cell function in HFD-fed mice. As shown in Fig. 2A-2B, serum glucagon and insulin levels were markedly elevated in HFD-fed mice compared with control, indicating compensatory hyperinsulinemia and dysregulated α-cell activity in HFD-fed mice. Interestingly, both 10 nM tirzepatide and the GIP analog [D-Ala²]-GIP normalized glucagon and insulin level in HFD-fed mice (Fig. 2A-2B). HOMA analyses revealed that only 10 nM tirzepatide enhanced β-cell function (HOMA-β; Fig. 2C), while 10 nM [D-Ala2]-GIP, 3 nM tirzepatide, and 10 nM tirzepatide all alleviated insulin resistance (HOMA-IR; Fig. 2D). In addition, 10 nM tirzepatide displayed more potency compared with 3 nM tirzepatide, indicating dose-dependent restoration of islet responsiveness for tirzepatide (Fig. 2A-2D).

**Figure 2.**
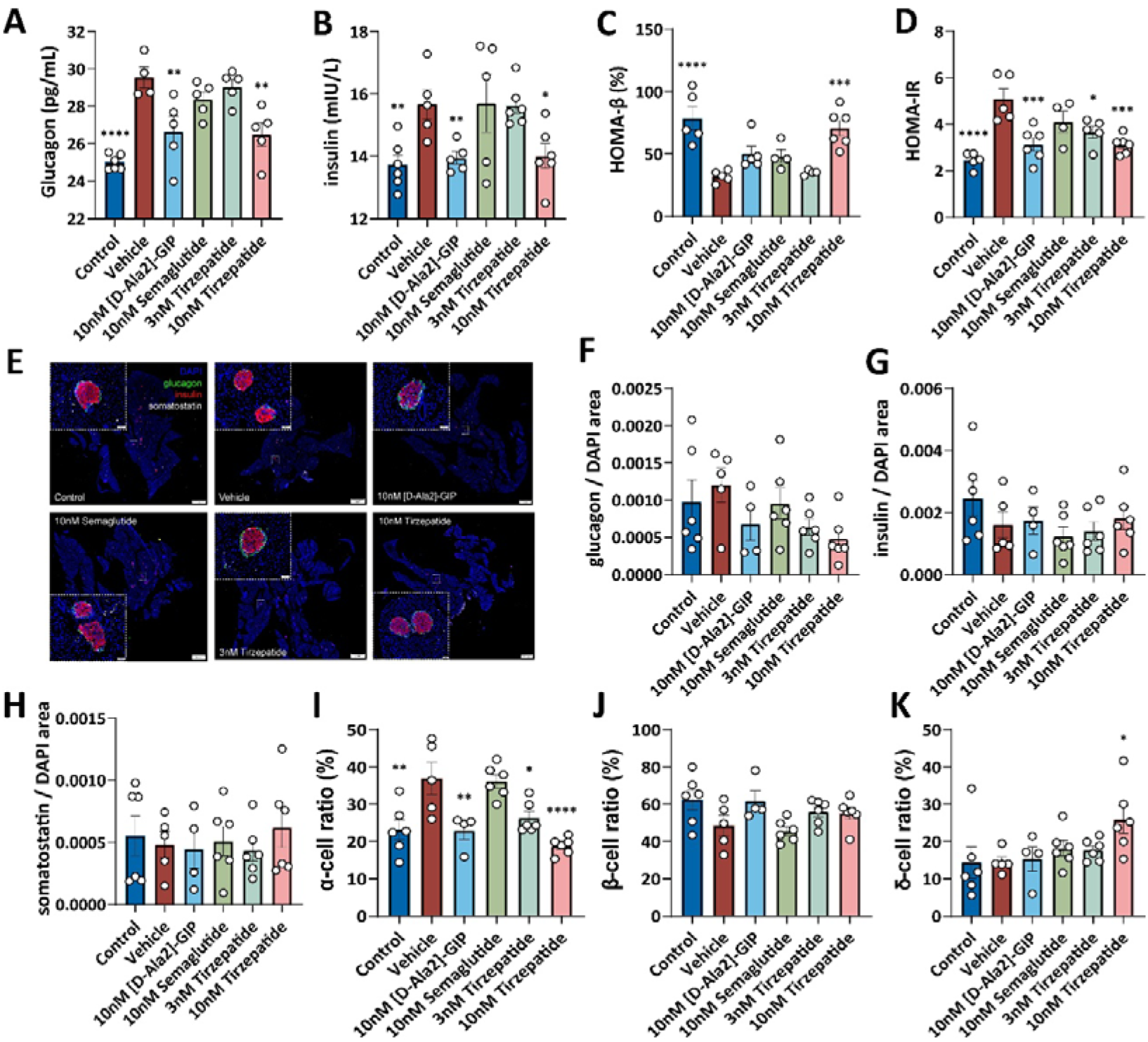
Tirzepatide improves β-cell function in HFD-fed mice. (**A-B**) Serum levels of glucagon (**A**) and insulin (**B**) in each group after different drug treatment, mice n ≥ 4. (**C-D**) HOMA-β (**C**) and HOMA-IR (**D**) indices in each group, mice n ≥ 4. (**E**) Representative images from whole-pancreas section immunofluorescence staining from each group, sections were stained with glucagon (green), insulin (red), somatostatin (white) and Dapi (blue). Scale bar indicates 1000 <u>μ</u>m and the small white dashed box was magnified in the corner. (**F-H**) Quantification of α cell (**F**), β cell (**G**) and δ (**H**) cell areas in each group based on whole-pancreas section immunofluorescence staining, n ≥ 4. (**I-K**) The ratio of α cell (**I**), β cell (**J**) and δ cell (**K**) among the three endocrine cell types within islets in each group, mice n ≥ 4. Data are presented as mean ± SEM, with significance determined by one-way ANOVA compared to the Vehicle group: \**p* < 0.05, \*\**p* < 0.01, \*\*\**p* < 0.001 and \*\*\*\**p* < 0.0001.

To explore how tirzepatide improves β-cell function, we followed to examine the endocrine cells mass and relative cell markers. We performed the whole-pancreas section immunostaining in these drugs treated mice. As shown in Fig. 2E-2H and Fig. S3, the islet number was not changed after the drugs administration, and the areas of α cell, β cell and δ cell also not affected. However, compared with the HFD group, the α cell ratio decreased the δ cell ratio increased and in 10 nM tirzepatide treated HFD mice, but not for β cell ratio (Fig. 2I-2K). Moreover, Ki67 and Caspase-3 immunostaining indicated that none of the treatments significantly affected cell proliferation or apoptosis (Fig. S4A-4D). These results suggest that the improved β-cell function was not attributable to changes in β-cell mass or proportion, but may involve regulation of β-cell identity.

Indeed, HFD feeding markedly increased the expression of ALDH1A3, which is a well-established marker of β-cell dedifferentiation [26], while all the drug treatments reversed the upregulation of ALDH1A3 in HFD-fed mice, indicating a rescue of β-cell dedifferentiation (Fig. 3A-3B). Importantly, HFD feeding led to a reduction in the level of PDX1 and MAFA, which are crucial transcription factors of β-cell identity and function (Fig. 3C-F). Whereas 3 nM tirzepatide and 10 nM tirzepatide treatment significantly restored the level of PDX1 and MAFA in β cells from HFD-fed mice (Fig. 3C-F). Taken together, these data demonstrate that tirzepatide reversed β-cell dedifferentiation and re-established β-cell identity, fostering a return to a mature and functional state in the setting of HFD-induced metabolic stress.

**Figure 3.**
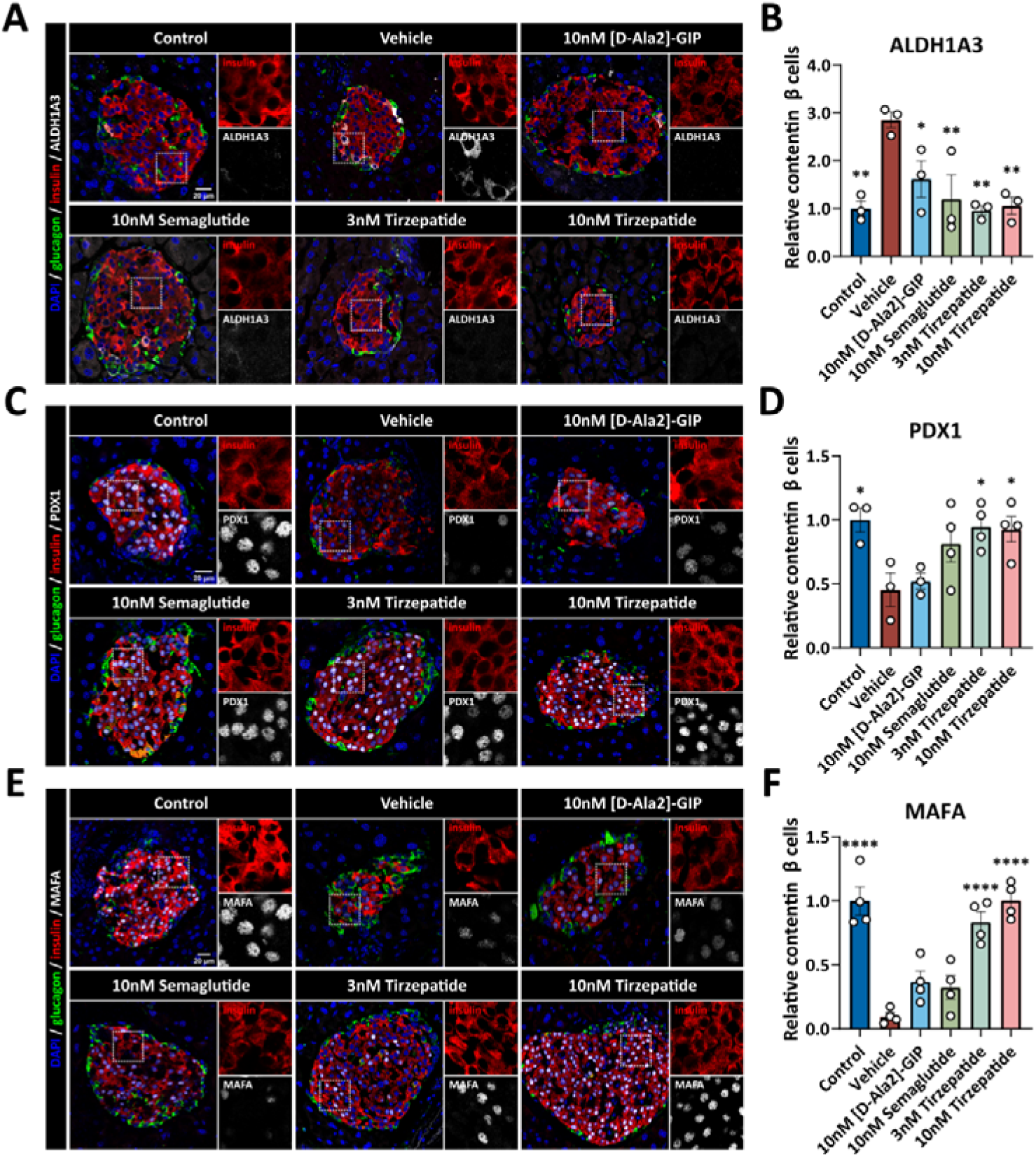
Tirzepatide restores the β-cell identity in HFD-fed mice. (**A**) Representative confocal images of islets by immunofluorescence staining from control group, HFD + vehicle, and HFD + 10 nM [D-Ala2]-GIP, HFD + 10nM semaglutide, HFD + 3 nM tirzepatide, HFD + 10 nM tirzepatide. Endocrine cells were labeled with insulin (red), glucagon (green), and ALDH1A3 (white). The nuclei were stained with Dapi (blue). The insets (dash box) of each group are magnified in the right panels of the images. (**B**) Quantification of relative levels of ALDH1A3 in each group, mice n ≥ 3. (**C**) Representative confocal images of islets by immunofluorescence staining from each group. Endocrine cells were labeled with insulin (red), glucagon (green), and PDX1 (white). The nuclei were stained with Dapi (blue). The insets (dash box) of each group are magnified in the right panels of the images. (**D**) Quantification of relative levels of PDX1 in each group, mice n ≥ 4. (**E**) Representative confocal images of islets by immuofluorescence staining from each group. Endocrine cells were labeled with insulin (red), glucagon (green), and MAFA (white). The nuclei were stained with Dapi (blue). The insets (dash box) of each group are magnified in the right panels of the images. Scale bar indicates 20 μm. (**F**) Quantification of relative levels of MAFA in each group, mice n ≥ 4. Data are presented as mean ± SEM, with significance determined by one-way ANOVA compared to the Vehicle group: \**p* < 0.05, \*\**p* < 0.01 and \*\*\*\**p* < 0.0001.

### 3.3 scRNA-seq identifies FOXO signaling is the key mediator for tirzepatide induced ***β***-cell function improvement

To elucidate the molecular mechanism underlying tirzepatide induced restoration of β-cell function, we performed single cell RNA sequencing (scRNA seq) on islets isolated from chow fed (CTR), HFD fed (HFD), and 10□nM tirzepatide treated (TZP) mice (Fig.□4A). After quality filtering and dimensional reduction, clustering analysis identified ten major pancreatic cell populations (Fig. 4B and Fig. S5A). Overall, the cell-type composition of islets remaining comparable across different groups (Fig. 4C-4D), suggesting that tirzepatide’s effects on islets from HFD-fed mice were improvement of endocrine cells function rather than composition of endocrine cells.

**Figure 4.**
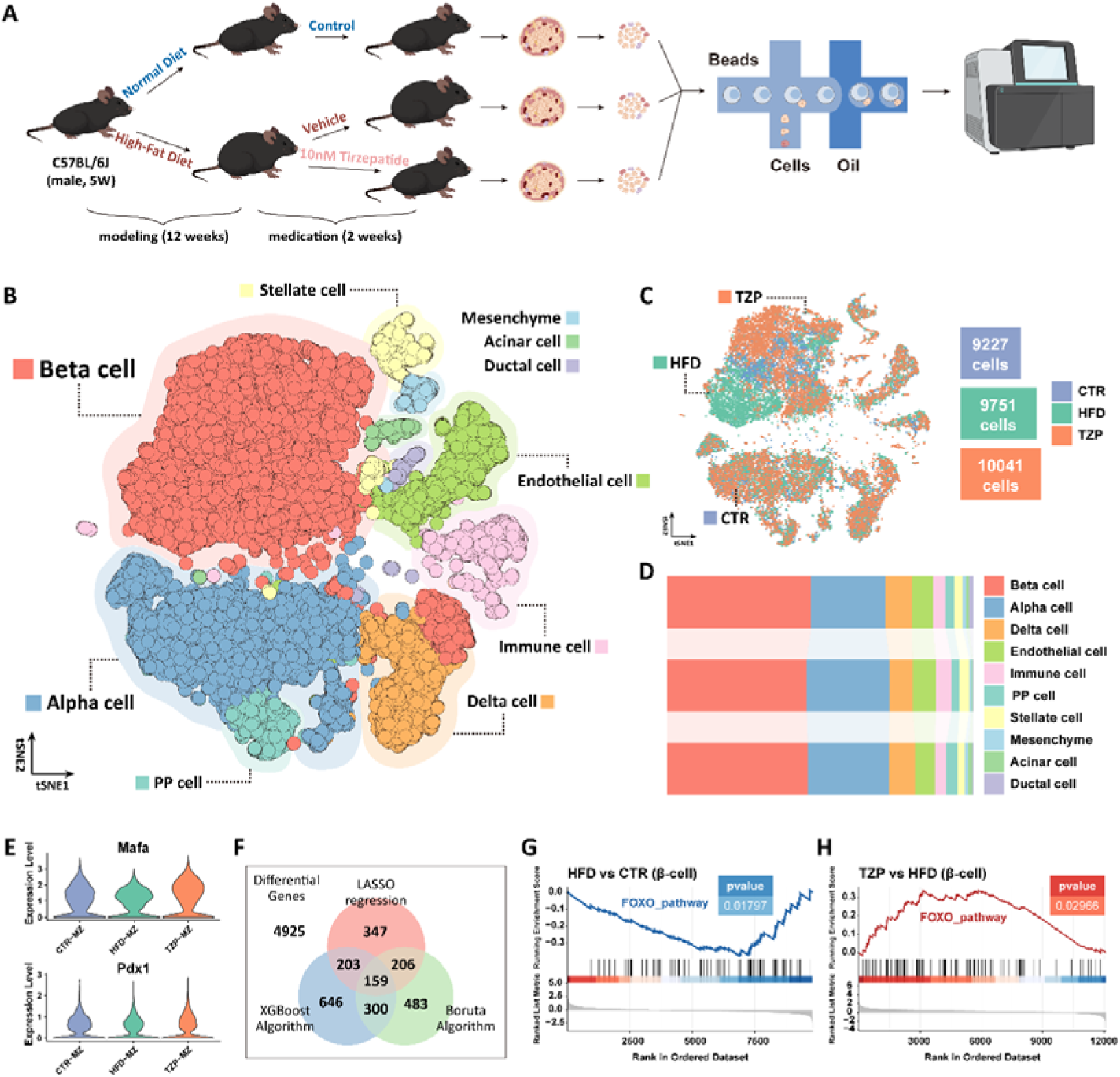
Single-cell RNA sequencing reveals tirzepatide improves β-cell function through the FOXO pathway. (**A**) Schematic workflow of single-cell RNA sequencing of mouse islets. (**B-C**) t-SNE plots showing all cells colored by cell type (**B**) and sample origin (**C**). (**D**) Proportions of different cell types among the three sample origins. (**E**) Expression levels of *Pdx1* and *Mafa* in β cells from the three origins. (**F**) Identification of common differentially expressed genes using three machine learning methods. (**G-H**) Gene set enrichment analysis (GSEA) of FOXO pathway related genes in β cells comparing the HFD and CTR groups (**G**), as well as the TZP and HFD groups (**H**).

We then isolated the β-cells from each group for further analysis (S5B-S5C). Within β-cells, HFD feeding led to reduction of *Pdx1* and *Mafa* expression, both of which were restored following tirzepatide treatment (Fig. 4E). To identify key molecules mediating this functional recovery, we first analyzed differentially expressed genes (DEGs) between the HFD and CTR groups, identifying 6,862 DEGs. Similarly, 8,201 DEGs were identified between the TZP and HFD groups. From these two comparisons, 4,925 overlapping DEGs were selected for further analysis. By machine-learning-based gene prioritization, included LASSO regression, Boruta algorithm and XGBoost algorithm, we derived a core set of 159 high-confidence DEGs through intersection analysis from these overlapped 4925 DEGs (Fig. 4F).

Subsequent KEGG (Fig. S6A) and protein protein interaction (PPI) (Fig. S6B) enrichment demonstrated a prominent enrichment of FOXO signaling pathway related genes, suggesting that both HFD feeding and tirzepatide treatment significantly modulate FOXO pathway activity in β cells. Moreover, the gene set enrichment analysis (GSEA) indicated FOXO pathway decreased in β cells from HFD-fed mice, but was restored in tirzepatide treated HFD-fed mice (Fig. 4G-H). Taken together, these findings indicate that tirzepatide enhances β-cell transcriptional programs through reactivation of the FOXO pathway, positioning it as a central mediator of β-cell function restoration.

### 3.4 Tirzepatide restores ***β***-cell function through upregulation of FOXO1

Given that FOXO1 serves as a central regulator of the FOXO pathway and plays a critical role in maintaining β-cell identity, oxidative defense, and insulin gene expression [6], we then examined the levels of FOXO1 and its phosphorylation state (p-FOXO1) in β-cell. Both total FOXO1 and p-FOXO1 were markedly reduced in β-cells from HFD-fed mice (Fig. 5A-D). However, 10 nM tirzepatide administration significantly restored the levels of both FOXO1 and p-FOXO1 (Fig. 5A-5D).

**Figure 5.**
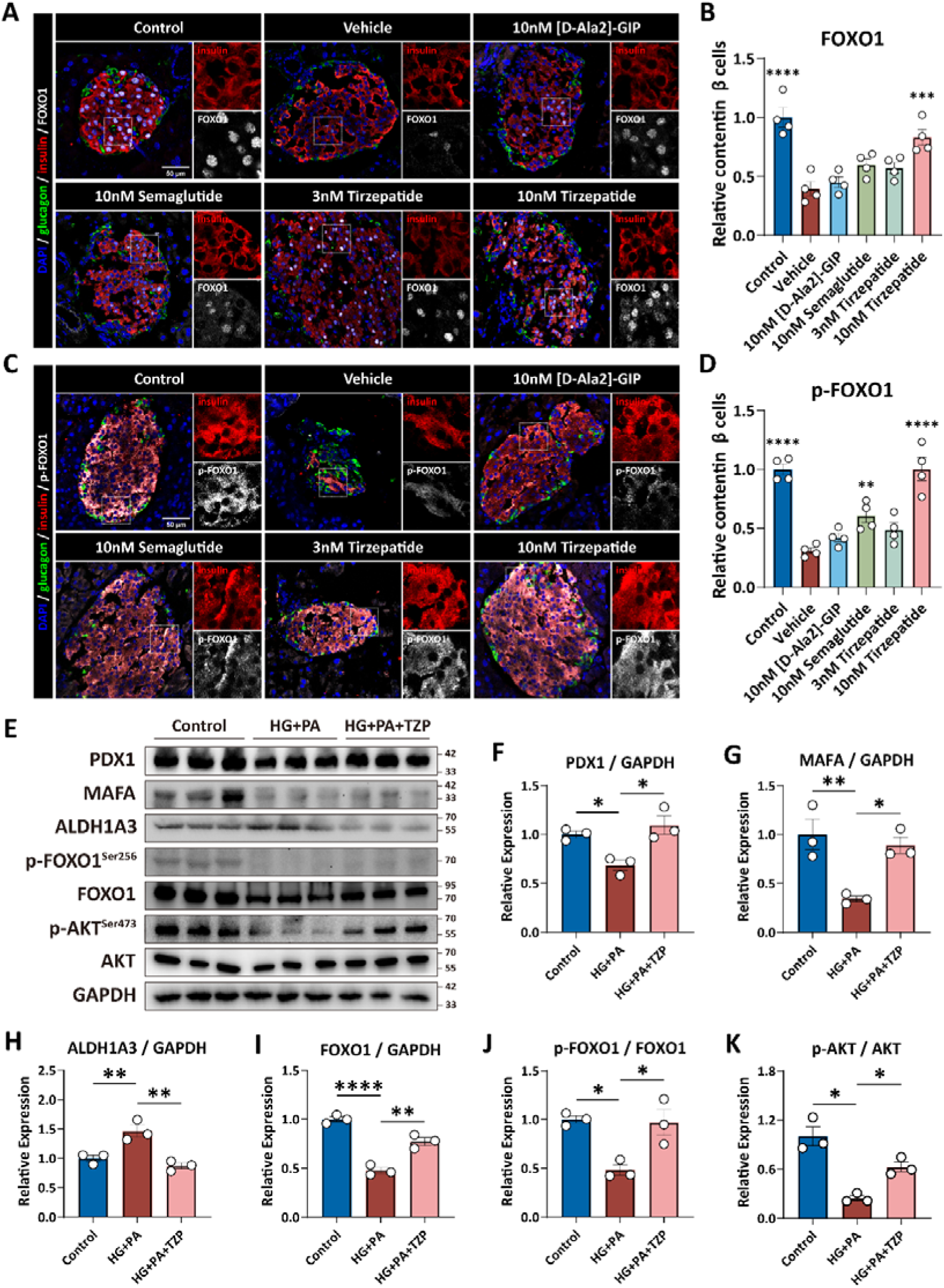
Tirzepatide restores β-cell function by upregulating FOXO1 expression in HFD-fed mice. (**A-B**) Representative images of FOXO1 staining in islet β cells (**A**) and quantification of relative levels (**B**) in each group. The insets (dash box) of each group are magnified in the right panels of the images. Scale bar indicates 50 μm. mice n ≥ 4 (**C-D**) Representative images of p-FOXO1 staining in islet β cells (**C**) and quantification of relative levels (**D**) in each group. The insets (dash box) of each group are magnified in the right panels of the images. Scale bar indicates 50 μm. mice n ≥ 4. (**E-L**) Western blot analysis of protein expression in MIN6 cells treated with high glucose and palmitate (HG+PA) or HG+PA plus tirzepatide (HG+PA+TZP) (**E**), with quantification of protein levels shown in (**F-L**). Data are presented as mean ± SEM. Statistical significance was determined by one-way ANOVA compared to the Vehicle group (**B** and **D**) and compared to the HG+PA group (**F-L**): \**p* < 0.05, \*\**p* < 0.01, \*\*\**p* < 0.001 and \*\*\*\**p* < 0.0001.

In MIN6 β cells, glucolipotoxic stress induced by high glucose plus palmitic acid (HG+PA) recapitulated key features of β cell dysfunction and dedifferentiation, as shown by decreased PDX1 and MAFA levels and increased ALDH1A3 expression (Fig. 5E-5K). However, tirzepatide treatment restored the levels of PDX1, MAFA and ALDH1A3 (Fig. 5E-5K). Consistent with the *in vivo* data, both FOXO1 (Fig. 5H) and p-FOXO1 levels decreased in HG+PA-treated MIN6 cells (Fig. 5E, 5I-5J), whereas tirzepatide treatment restored their levels (Fig. 5I-5J). Moreover, as the primary upstream cascade of FOXO1, AKT and its activated form (p-AKT) both decreased in HG+PA-treated MIN6 cells and restored after tirzepatide treatment (Fig. 5E, 5K).

Together, these results demonstrated that both HFD feeding and glucolipotoxicity down regulate FOXO1 levels in β-cells, whereas tirzepatide treatment reverses this suppression. Restoration of FOXO1 level and activity may underlie the recovery of PDX1 and MAFA, and overall β-cell functional capacity following tirzepatide administration.

### 3.5 Tirzepatide improves pancreatic ***β***-cell function and insulin sensitivity in patients with type 2 diabetes

Finally, we assessed the therapeutic efficacy of tirzepatide in patients with type 2 diabetes. Thirteen clinically diagnosed T2DM patients received intravenous tirzepatide with cumulative doses of 30-60 mg in three months therapeutical treatment (maximum single dose 7.5 mg; Fig. 6A). The detail demographic and clinical characteristics of these patients are provided in Supplementary Table 1.

**Figure 6.**
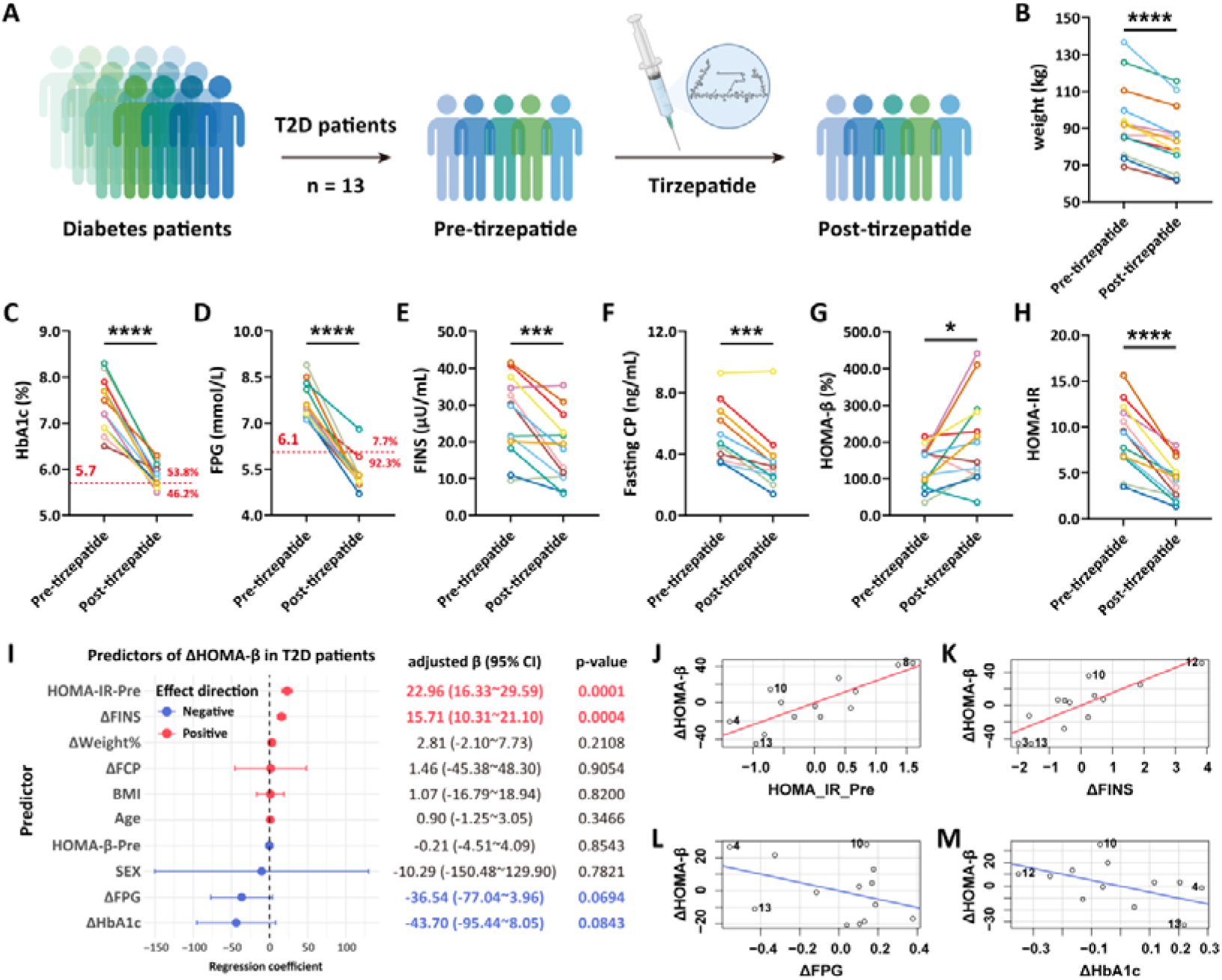
Tirzepatide improves β-cell function and insulin sensitivity in T2DM patients. (**A**) Assessment of 13 clinically diagnosed T2DM patients before and after tirzepatide treatment. (**B-H**) Changes in body weight (**B**) and key metabolic parameters (**C**-**H**) following three months of tirzepatide administration: HbA1c (**C**), fasting plasma glucose (**D**), fasting insulin (**E**), fasting C-peptide (**F**), HOMA-β (**G**), and HOMA-IR (**H**). (**I**) Multivariable regression analysis identifying independent predictors associated with HOMA-β improvement (ΔHOMA-β, defined as post-treatment HOMA-β minus pre-treatment HOMA-β). (**J-M**) Baseline HOMA-IR (HOMA_IR_Pre, **J**) and the reduction in fasting insulin (ΔFINS, **K**), fasting plasma glucose (ΔFPG, **L**) and HbA1c (ΔHbA1c, **M**) were independently correlated with the extent of HOMA-β improvement. Statistical significance was determined by paired Student’s t-test: \**p* < 0.05, \*\*\**p* < 0.001 and \*\*\*\**p* < 0.0001.

Body weight, glycated hemoglobin A1c (HbA1c), fasting plasma glucose (FPG), fasting insulin (FINS), and fasting C-peptide (FC) were measured in patients both before (pre-tirzepatide group) and after (post-tirzepatide group) treatment. Tirzepatide markedly reduced body weight (Fig. 6B) and significantly improved glycemic control in T2D patients, with 46.2% of patients achieving normal HbA1c levels after three months therapy (Fig. 6C) and 92.3% attaining normalized fasting glucose (Fig. 6D). Both fasting insulin and C-peptide levels declined after treatment (Fig. 6E-6F), indicating reduced insulin demand in these patients. Moreover, the HOMA-β increased and the HOMA-IR decreased in T2D patients after treatment, which suggested tirzepatide therapy improves β-cell function and attenuated insulin resistance in in T2D patients (Fig. 6G-6H). The changes of tirzepatide induced β-cell function and insulin resistance were consistent with observation in diabetic mice (Fig. 2C-2D).

To identify parameters associated with β-cell function improvement, we performed a multivariable logistic regression analysis using the change in HOMA-β (ΔHOMA-β; defined as post-treatment minus pre-treatment, with Δ for all subsequent variables similarly indicating post-treatment minus pre-treatment) as the dependent variable. As shown in Fig. 6, baseline HOMA IR (HOMA IR Pre, Fig. 6I and 6J) and ΔFINS (Fig. 6I and 6K) displayed potential positive correlations with ΔHOMA-β, suggesting that in this cohort of tirzepatide-treated T2D patients, those with higher baseline insulin resistance tended to exhibit greater β-cell functional recovery. And patients with more pronounced β-cell improvement experienced smaller decreases in circulating insulin levels. Additionally, although not statistically significant, ΔFPG and ΔHbA1c displayed negative trends with ΔHOMA β (Fig.□6I, 6L and 6M), indicating that larger glucose lowering responses generally accompanied greater β cell functional restoration.

Taken together, these results demonstrate that tirzepatide not only improves β-cell function but also reduces insulin resistance in T2D patients, suggested that tirzepatide represents a therapeutic strategy is able to addresses β-cell dysfunction in T2DM.

## 4. DISCUSSION

In this study, we demonstrated that tirzepatide exerts profound benefits in β-cell function improvement, both in HFD-induced T2D mouse model and T2D patients. In HFD-induced T2D mouse model, we show that tirzepatide not only ameliorates systemic metabolic dysfunction, including obesity, hepatic steatosis, dyslipidemia, and insulin resistance, but also restores β-cell identity and functional competence without increasing β-cell mass. These preclinical findings are supported by results from a clinical cohort, in which tirzepatide treatment improved glycemic control, reduced insulin demand, and increased HOMA-β while lowering HOMA IR, underscoring the translational relevance of our observations.

Mechanistically, we reveal that tirzepatide restores β-cell function by rescuing FOXO1 signaling, which in turn up regulates the expression of PDX1 and MAFA, thereby reversing β-cell dedifferentiation and re-establishing β cell identity. Consistent with previous studies, our results indicated that HFD feeding and glucolipotoxicity reduce FOXO1 levels, suppress PDX1 and MAFA expression, and increase ALDH1A3, indicated β-cell dedifferentiation [6, 7, 27, 28] (Fig. 3–Fig. 5). Tirzepatide administration restores both total and phosphorylated FOXO1 levels, and increased PDX1 and MAFA levels, and improves insulin secretory capacity (Fig. 3–Fig. 5). Moreover, *in vitro* glucolipotoxicity assays further confirmed that the AKT pathway was suppressed under metabolic stress, leading to reduced p-FOXO1 in β cells. While tirzepatide incubation reactivates the AKT pathway (Fig. 5E and 5K). These results are in line with previous studies implicating FOXO1 as a master regulator of β-cell plasticity under metabolic stress [6, 29, 30]. Most importantly, these results may indicate that dual incretin receptor activation can reverse stress-induced β-cell dysfunction through AKT-FOXO1-PDX1/MAFA axis.

Other investigators have likewise reported that tirzepatide enhances β-cell function and insulin sensitivity across diverse T2D populations [13, 15, 16]. Partially consistent with some of these findings, our study shows that improvements in HOMA-β and HOMA-IR correlate with baseline insulin resistance and reductions in HbA1c in T2D patients (Fig. 6). We further revealed that improvements in HOMA-β positively correlate with baseline insulin resistance and reductions in HbA1c, but not with age, suggesting that metabolic context rather than chronological age determines β-cell recovery potential.

Notably, the limited effects of semaglutide on β-cell functional recovery observed in our HFD-fed mice differ from several previous reports. Semaglutide, as well as other GLP-1 receptor agonists, has been shown to improve glycemic homeostasis in T2D mouse models and to restore the expression of key β-cell transcription factors [31, 32]. The discrepancy between these findings and our results is likely multifactorial. Firstly, dosing differences may represent a critical determinant of therapeutic efficacy. Previous studies use 40 μg/kg semaglutide [31, 32], or 200 nmol/kg semaglutide for administration [33], while we use 10 nM semaglutide. Secondly, differences in the choice of T2D mouse models and disease induction strategies may further account for the inconsistent observations. Previous studies use db/db mice [33], or 4 weeks high-fat diet-fed plus streptozotocin (STZ) mice [31], we used 12 high-fat diet-fed mice. Together, these considerations suggest that the impact of semaglutide on β-cell recovery is highly context dependent, being influenced by disease stage, dosing regimen, and experimental model selection.

In summary, our findings provide partial but compelling evidence that tirzepatide acts through a dual mechanism. First, it improves systemic metabolism by reducing body weight, redistributing lipids, and enhancing insulin sensitivity. Second, it directly restores β-cell transcriptional and functional competence through reactivation of the FOXO1-PDX1/MAFA axis. This dual mode of action positions tirzepatide as a rational therapeutic strategy that targets both the causes and consequences of β-cell dysfunction in T2DM.

Nevertheless, our study also has some limitations. The sample size of patient is relatively small, and more comprehensive clinical characterization will require validation in larger cohorts. Moreover, the detail mechanism of how tirzepatide regulates FOXO1 for regulation of β-cell functional genes need to be further investigated. Furthermore, how tirzepatide regulates the GLP1R/GIPR dual receptors also need to be explored.

## Supporting information

All supplemantal data

## Acknowledgements

We thank members of the Li lab for constructive discussions. We also thank Zhen Li, Hong Yun, Yuhong Chen, Shan Jiang from School of Pharmaceutical Sciences, Xiamen University; Lei Huang from School of Life Sciences, Xiamen University, for technical support.

## Author Contributions

M.L. and N.C. supervised this work, and, as such, had full access to all of the data in the study and take responsibility for the integrity of the data and the accuracy of the data analysis. M.L. and N.C. designed the study. Z.L., J.G. and Y.C. performed key experiments. M.L., N.C., Z.L., J.G., Y.C., T.Z., X.L., S.Z., Q.R., and Z.W. participated in the planning of the work and the interpretation of the results. Z.L. drafted the manuscript. M.L. and N.C. revised the paper.

## Data Availability

The authors declare that the data supporting the findings of this study are available within the Article, and its Supplementary Information. The original scRNA-seq data generated in this study have been deposited in the BioProject database under accession code PRJNA1402239 [https://dataview.ncbi.nlm.nih.gov/object/PRJNA1402239?reviewer=hlfnqkftqcmedq 6ctdapdkalql]. Source data are provided with this paper.

## Abbreviations

T2D: type 2 diabetes (T2D)
HFD: High-fat diet
GLP-1: Glucagon-like peptide-1
GIP: Glucose dependent insulinotropic polypeptide
FOXO1: Forkhead box protein O1
PDX1: Pancreatic duodenum homeobox-1
MAFA: MAF bZIP transcription factor A
ALDH1A3: Aldehyde dehydrogenase 1A3
HOMA-β: Homeostasis model assessment of β-cell function
HOMA-IR: Homeostasis model assessment of insulin resistance
HbA1c: Hemoglobin A1C

## References

[1] Butler AE, Janson J, Bonner-Weir S, Ritzel R, Rizza RA, Butler PC (2003) Beta-cell deficit and increased beta-cell apoptosis in humans with type 2 diabetes. Diabetes 52(1): 102–110. 10.2337/diabetes.52.1.102

[2] Reaven GM (1988) Banting lecture 1988. Role of insulin resistance in human disease. Diabetes 37(12): 1595–1607. 10.2337/diab.37.12.1595

[3] Accili D, Deng Z, Liu Q (2025) Insulin resistance in type 2 diabetes mellitus. Nat Rev Endocrinol 21(7): 413–426. 10.1038/s41574-025-01114-y

[4] Eguchi K, Nagai R (2017) Islet inflammation in type 2 diabetes and physiology. J Clin Invest 127(1): 14–23. 10.1172/JCI88877

[5] Poitout V, Robertson RP (2008) Glucolipotoxicity: fuel excess and beta-cell dysfunction. Endocr Rev 29(3): 351–366. 10.1210/er.2007-0023

[6] Talchai C, Xuan S, Lin HV, Sussel L, Accili D (2012) Pancreatic beta cell dedifferentiation as a mechanism of diabetic beta cell failure. Cell 150(6): 1223–1234. 10.1016/j.cell.2012.07.029

[7] Guo S, Dai C, Guo M, et al. (2013) Inactivation of specific beta cell transcription factors in type 2 diabetes. J Clin Invest 123(8): 3305–3316. 10.1172/JCI65390

[8] Rohm TV, Meier DT, Olefsky JM, Donath MY (2022) Inflammation in obesity, diabetes, and related disorders. Immunity 55(1): 31–55. 10.1016/j.immuni.2021.12.013

[9] Drucker DJ, Holst JJ (2023) The expanding incretin universe: from basic biology to clinical translation. Diabetologia 66(10): 1765–1779. 10.1007/s00125-023-05906-7

[10] Campbell JE, Muller TD, Finan B, DiMarchi RD, Tschop MH, D’Alessio DA (2023) GIPR/GLP-1R dual agonist therapies for diabetes and weight loss-chemistry, physiology, and clinical applications. Cell Metab 35(9): 1519–1529. 10.1016/j.cmet.2023.07.010

[11] Marso SP, Bain SC, Consoli A, et al. (2016) Semaglutide and Cardiovascular Outcomes in Patients with Type 2 Diabetes. N Engl J Med 375(19): 1834–1844. 10.1056/NEJMoa1607141

[12] Pratley RE, Aroda VR, Lingvay I, et al. (2018) Semaglutide versus dulaglutide once weekly in patients with type 2 diabetes (SUSTAIN 7): a randomised, open-label, phase 3b trial. Lancet Diabetes Endocrinol 6(4): 275–286. 10.1016/S2213-8587(18)30024-X

[13] Frias JP, Nauck MA, Van J, et al. (2018) Efficacy and safety of LY3298176, a novel dual GIP and GLP-1 receptor agonist, in patients with type 2 diabetes: a randomised, placebo-controlled and active comparator-controlled phase 2 trial. Lancet 392(10160): 2180–2193. 10.1016/S0140-6736(18)32260-8

[14] Jastreboff AM, Aronne LJ, Ahmad NN, et al. (2022) Tirzepatide Once Weekly for the Treatment of Obesity. N Engl J Med 387(3): 205–216. 10.1056/NEJMoa2206038

[15] Frias JP, Davies MJ, Rosenstock J, et al. (2021) Tirzepatide versus Semaglutide Once Weekly in Patients with Type 2 Diabetes. N Engl J Med 385(6): 503–515. 10.1056/NEJMoa2107519

[16] Frias JP, De Block C, Brown K, et al. (2024) Tirzepatide Improved Markers of Islet Cell Function and Insulin Sensitivity in People With T2D (SURPASS-2). J Clin Endocrinol Metab 109(7): 1745–1753. 10.1210/clinem/dgae038

[17] Winzell MS, Ahrén B (2004) The high-fat diet-fed mouse: a model for studying mechanisms and treatment of impaired glucose tolerance and type 2 diabetes. Diabetes 53 Suppl 3: S215–219. 10.2337/diabetes.53.suppl_3.s215

[18] Stuart T, Butler A, Hoffman P, et al. (2019) Comprehensive Integration of Single-Cell Data. Cell 177(7): 1888–1902.e1821. 10.1016/j.cell.2019.05.031

[19] Ritchie ME, Phipson B, Wu D, et al. (2015) limma powers differential expression analyses for RNA-sequencing and microarray studies. Nucleic Acids Res 43(7): e47. 10.1093/nar/gkv007

[20] Ogata H, Goto S, Sato K, Fujibuchi W, Bono H, Kanehisa M (1999) KEGG: Kyoto Encyclopedia of Genes and Genomes. Nucleic Acids Res 27(1): 29–34. 10.1093/nar/27.1.29

[21] Friedman J, Hastie T, Tibshirani R (2010) Regularization Paths for Generalized Linear Models via Coordinate Descent. J Stat Softw 33(1): 1–22

[22] Kursa MB, Rudnicki WR (2010) Feature Selection with the Boruta Package. Journal of Statistical Software 36(11): 1–13

[23] Chen T, Guestrin C (2016) XGBoost: A Scalable Tree Boosting System. In, pp 785–794

[24] Viberti G, Kahn SE, Greene DA, et al. (2002) A diabetes outcome progression trial (ADOPT): an international multicenter study of the comparative efficacy of rosiglitazone, glyburide, and metformin in recently diagnosed type 2 diabetes. Diabetes Care 25(10): 1737–1743. 10.2337/diacare.25.10.1737

[25] Gu Z, Gu L, Eils R, Schlesner M, Brors B (2014) circlize Implements and enhances circular visualization in R. Bioinformatics 30(19): 2811–2812. 10.1093/bioinformatics/btu393

[26] Kim-Muller JY, Fan J, Kim YJ, et al. (2016) Aldehyde dehydrogenase 1a3 defines a subset of failing pancreatic β cells in diabetic mice. Nat Commun 7: 12631. 10.1038/ncomms12631

[27] Kim-Muller JY, Fan J, Kim YJ, et al. (2016) Aldehyde dehydrogenase 1a3 defines a subset of failing pancreatic beta cells in diabetic mice. Nat Commun 7: 12631. 10.1038/ncomms12631

[28] Wang LK, Kong CC, Yu TY, et al. (2025) Endoplasmic reticulum stress and forkhead box protein O1 inhibition mediate palmitic acid and high glucose-induced beta-cell dedifferentiation. World J Diabetes 16(5): 95431. 10.4239/wjd.v16.i5.95431

[29] Liu N, Li R, Cao J, et al. (2023) The inhibition of FKBP5 protects beta-cell survival under inflammation stress via AKT/FOXO1 signaling. Cell Death Discov 9(1): 247. 10.1038/s41420-023-01506-x

[30] Habibe JJ, Clemente-Olivo MP, Scheithauer TPM, et al. (2022) Glucose-mediated insulin secretion is improved in FHL2-deficient mice and elevated FHL2 expression in humans is associated with type 2 diabetes. Diabetologia 65(10): 1721–1733. 10.1007/s00125-022-05750-1

[31] Luo Y, Li JE, Zeng H, Zhang Y, Yang S, Liu J (2025) Semaglutide alleviates the pancreatic beta cell function via the METTL14 signaling and modulating gut microbiota in type 2 diabetes mellitus mice. Life Sci 361: 123328. 10.1016/j.lfs.2024.123328

[32] Marinho TS, Martins FF, Cardoso LEM, Aguila MB, Mandarim-de-Lacerda CA (2022) Pancreatic islet cells disarray, apoptosis, and proliferation in obese mice. The role of Semaglutide treatment. Biochimie 193: 126–136. 10.1016/j.biochi.2021.10.017

[33] Iwamoto Y, Kimura T, Dan K, et al. (2024) Tirzepatide, a dual glucose-dependent insulinotropic polypeptide/glucagon-like peptide 1 receptor agonist, exhibits favourable effects on pancreatic beta-cells and hepatic steatosis in obese type 2 diabetic db/db mice. Diabetes Obes Metab 26(12): 5982–5994. 10.1111/dom.15972

