## Supplementary material for "Tirzepatide improves pancreatic β-cell function in mice and patients with type 2 diabetes": All supplemantal data

**Supplemental Figure**

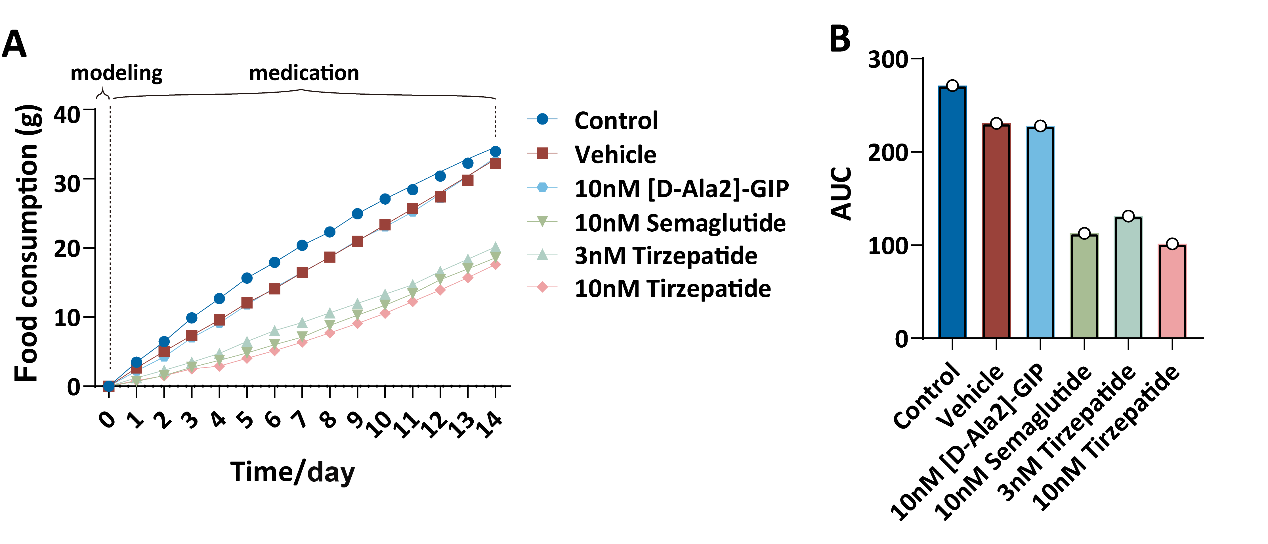

**Figure S1. Food intake analysis in each group of mice.**

(**A-B**) Food intake dynamics in each group (**A**) and corresponding AUC analysis (**B**).

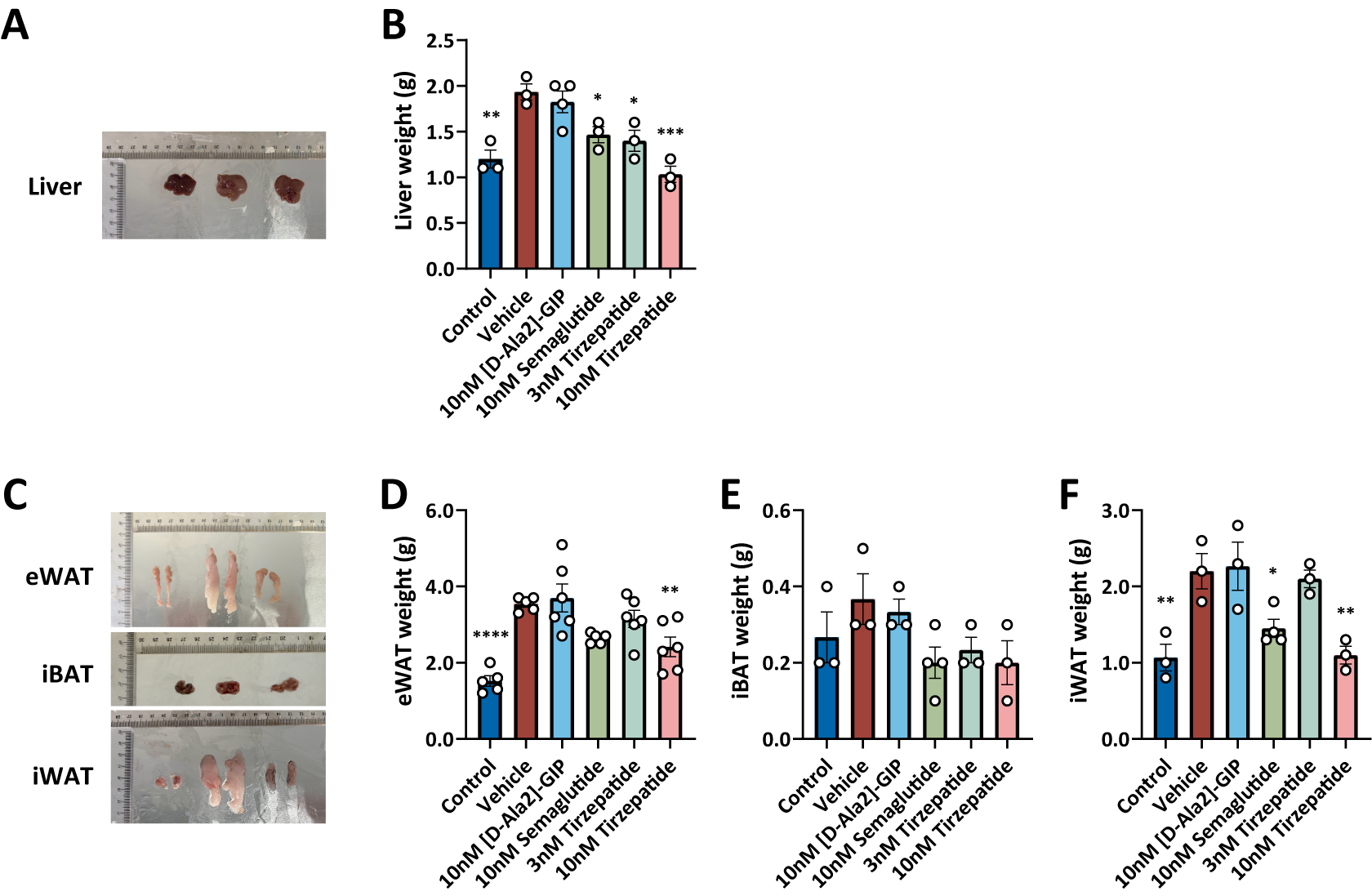

**Figure S2. Liver and adipose tissue weight measurements in each group of mice.**

(**A-B**) Representative images of mouse livers (**A**) and corresponding liver weight measurements (**B**), mice n ≥ 3. (**C-F**) Representative images of eWAT, iBAT, and iWAT (**C**) with corresponding weight measurements (**D-F**), mice n ≥ 3. Data are presented as mean ± SEM, with significance determined by one-way ANOVA compared to the Vehicle group: **p* < 0.05, ***p* < 0.01, ****p* < 0.001 and *****p* < 0.0001.

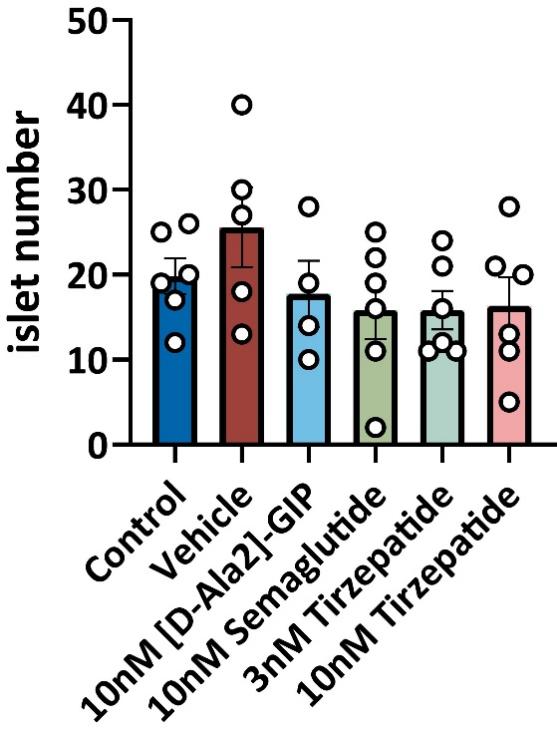

**Figure S3. Pancreatic islets counts in each group of mice.**

Quantification of pancreatic islet counts in each group of mice, mice n ≥ 4. Data are presented as mean ± SEM. Statistical significance was determined by one-way ANOVA compared to the Vehicle group.

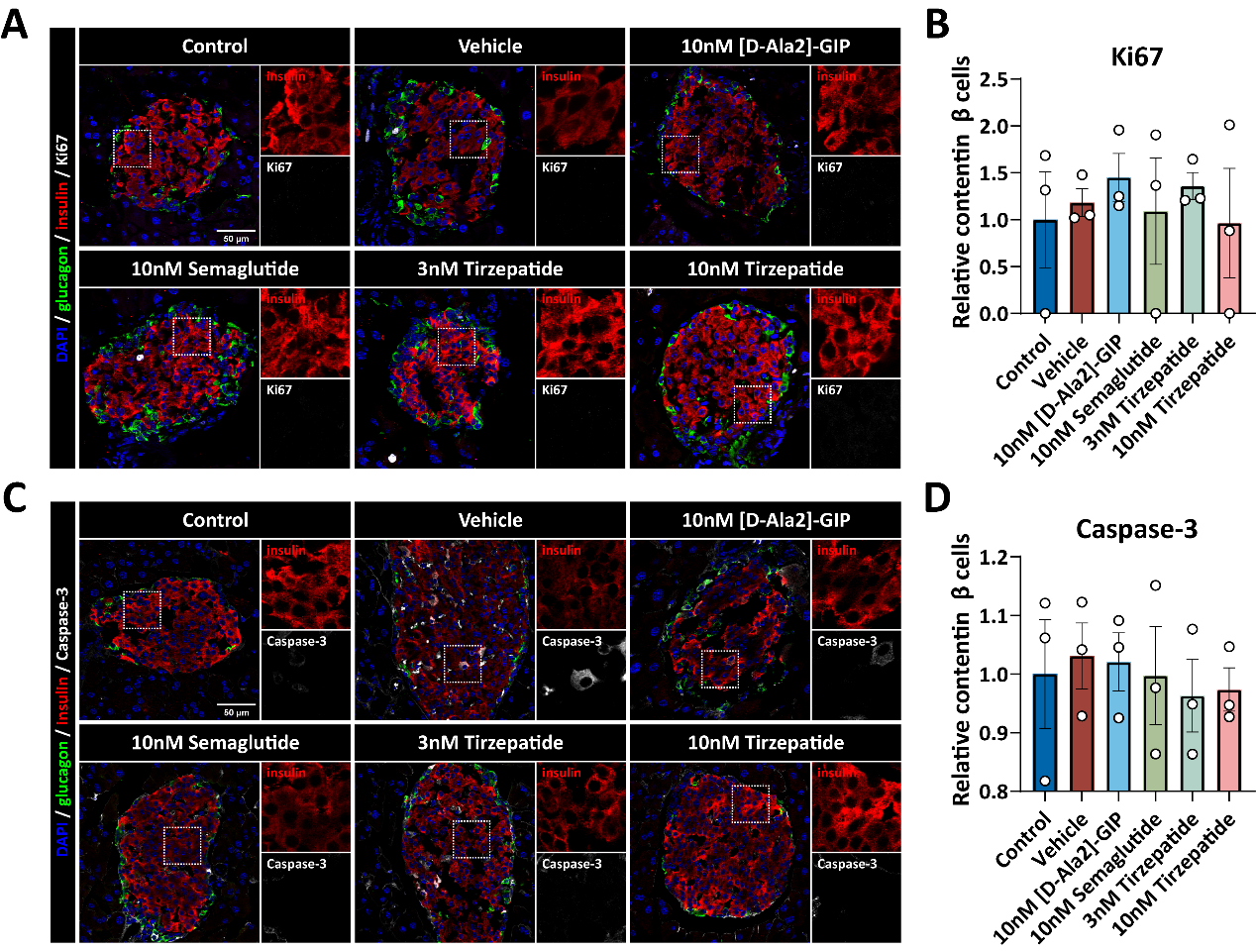

**Figure S4. Detection of β-cell Ki67 and Caspase-3 levels in each group of mice.**

(**A-B**) Representative images of Ki67 staining in islet β cells (**A**) and quantification of relative levels (**B**) in each group, mice n ≥ 3. Scale bar indicates 50 μm. (**C-D**) Representative images of Caspase-3 staining in islet β cells (**C**) and quantification of relative levels (**D**) in each group, mice n ≥ 3. Statistical significance was determined by one-way ANOVA compared to the Vehicle group.

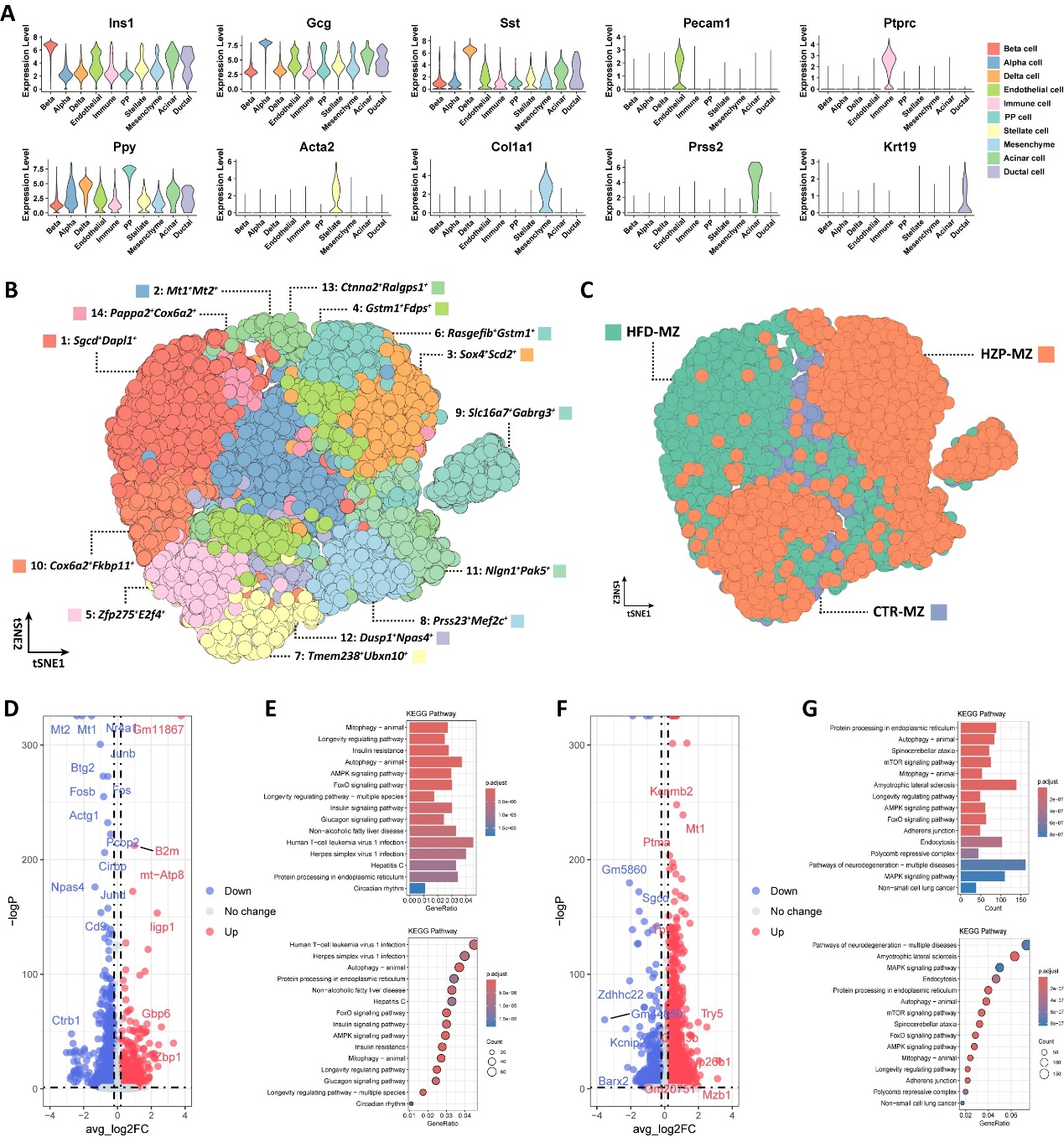

**Figure S5. Tirzepatide modulates the β-cell *Foxo* pathway in HFD-fed mice.**

(**A**) Violin plots showing expression of marker genes for each cell type. (**B-C**) t-SNE plots of all β cells colored by subtype (**B**) and sample origin (**C**). (**D-E**) Volcano plot of DEGs between HFD- and CTR-derived β cells (**D**) and KEGG pathway enrichment of these DEGs (**E**). (**F-G**) Volcano plot of DEGs between TZP- and HFD-derived β cells (**F**) and KEGG pathway enrichment of these DEGs (**G**).
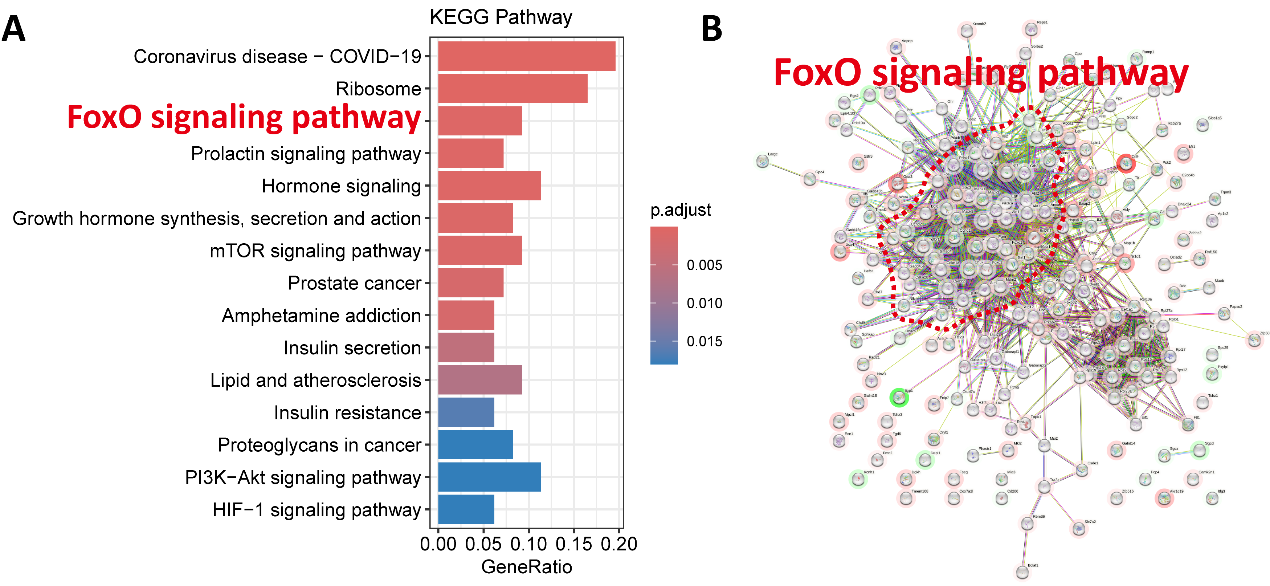

**Figure S6. FOXO pathway enrichment of genes identified by machine learning.**

KEGG pathway analysis (**A**) and protein-protein interaction (PPI) network analysis (**B**) demonstrated that genes selected by machine learning were significantly enriched in the FOXO signaling pathway.
